# Pathogenesis of Zika Virus Infection via Rectal Route

**DOI:** 10.1101/128876

**Authors:** Laura E. Martínez, Gustavo Garcia, Deisy Contreras, Danyang Gong, Ren Sun, Vaithilingaraja Arumugaswami

## Introduction

Zika virus (ZIKV) is a mosquito-borne flavivirus originally confined to Africa and Asia that has spread to islands located in Southeast Asia, and most recently to the Americas and the Caribbean. Approximately 80% of infected individuals are asymptomatic, while the remaining infected population exhibit mild febrile syndrome such as rash, conjunctivitis, and arthralgia. In some adults, ZIKV causes neurotropic Guillain–Barré syndrome^1^. Vertical transmission of ZIKV in infected mothers causes fetal growth restriction, microcephaly, and congenital eye disease^2-5^. Cases of ZIKV sexual transmission from male to female^6-8^, male to male^9^, and a suspected case of female to male transmission^10^ have been reported. ZIKV has been detected in the semen of infected males^11-15^, even after months of symptom onset^16-19^. Viral persistence in the testes and semen can increase the risk of ZIKV transmission through rectal route in men having sex with men (MSM) and between heterosexual partners. The anorectal mucosa is a major entry site for HIV-1 transmission among MSM^20^, and for the acquisition and transmission of other sexually transmitted diseases, such as syphilis, chlamydia, and gonorrhea. Although the risk of ZIKV acquisition through the rectal route is high, no pathobiological information is available. Here, we describe the establishment of a rectal route of ZIKV infection system using immunocompromised (*ifnar1*^*-/-*^) male mice to determine their susceptibility to ZIKV and to assess viral dissemination to male reproductive organs. We found that rectal inoculation of ZIKV results in viremia with non-lethal infection. The rectal mucosa is susceptible to ZIKV entry and replication. Following rectal inoculation, ZIKV establishes active testicular infection that persists at least 21 days. During the acute phase of infection, the highest viral load was observed in the spleen, with inflammatory and immune cellular infiltration. Macrophages in the splenic red pulp are the target cells for ZIKV infection.

Mouse models of ZIKV pathogenesis have shown that mice deficient in type I IFN receptor (*ifnar1*^*-/-*^ or A129) signaling pathway develop neurological disease in adults and congenital infection in pregnant females^21-24^. Adult immunocompetent (wild-type) mice are resistant to ZIKV infection due to a robust innate immune response that limits infection and spread. Thus, the *ifnar1*^*-/-*^ mouse model has become a widely used *in vivo* system to investigate the pathogenesis of ZIKV diseases. Subcutaneous or intravaginal ZIKV infection of *ifnar1*^*-/-*^ female mice leads to the activation of systemic and localized immune responses and the establishment of congenital and neurological diseases^23^,^25^,^26^. Vaginal exposure of ZIKV during the first trimester of pregnancy leads to fetal growth restriction and brain infection in wild-type mice, and loss of pregnancy in *ifnar1*^-/-^ mice^26^. In *ifnar1*^*-/-*^ male mice, ZIKV robustly infects the brain, spinal cord, and testes^22^. ZIKV causes injury in the testes and epididymis of male mice, further reducing the levels of testosterone and oligospermia^27^. Moreover, ZIKV persists within the testes of *ifnar1*^-/-^ mice following subcutaneous inoculation and causes testicular atrophy at 21 days post-infection^28^. ZIKV-mediated damage to the testes may lead to male infertility^29^. Recently, it was shown that intraperitoneally infected AG129 (interferon α/β and -γ receptor knockout) male mice had persistent testicular infection for more than a month, and the semen contained infectious ZIKV from 1 to 3 weeks post-inoculation^30^. In addition, ZIKV infection was documented in 50% of female mice mated to infected non-vasectomized male mice^30^. Collectively, these findings provide important insights on the long-term potential effects of ZIKV infection of female and male reproductive organs. Currently, there is no knowledge regarding ZIKV acquisition via rectal route of infection. Further studies are needed to understand ZIKV replication and persistence within different tissue compartments after sexual transmission, including the rectal mucosa.

In this study, we used *ifnar1*^-/-^ mice to study the pathogenesis of ZIKV Asian genotype strain PRVABC59 in 4-6-week-old male mice by rectal inoculation (3.4×10^6^ pfu per mouse), and followed disease progression for 21 days post-inoculation (dpi) (Fig. 1A). Mice were closely observed twice a day during the experimental period. Weight loss and neurological signs were monitored at various timepoints. We found that rectal inoculation of ZIKV does not cause mortality in *ifnar1*^-/-^ male mice (Fig. 1B). Two of five infected mice showed lethargy and transient sickness, with relative body weight loss (12-14%) at 10 dpi (Fig. 1C). These mice were closely monitored and they steadily recovered by 21 dpi (Fig. 1C). In addition, infected mice showed peak viremia at 7 dpi (Fig. 1D). In an independent experiment, we tested the subcutaneous model of infection with PRVABC59 (2×10^6^ pfu/mouse) to confirm its pathogenesis and development of lethal disease. Consistent with previous reports using A129 mice^31^, subcutaneously infected mice demonstrated significant weight loss and eventually succumbed to infection (Supplementary Fig. 1A-C). Infected mice had high acute viremia as early as 3 dpi and exhibited malaise, incoordination and neurological signs of posterior hind leg paralysis by 6 -7 dpi (Supplementary Fig. 1C-D).

**Figure 1.**
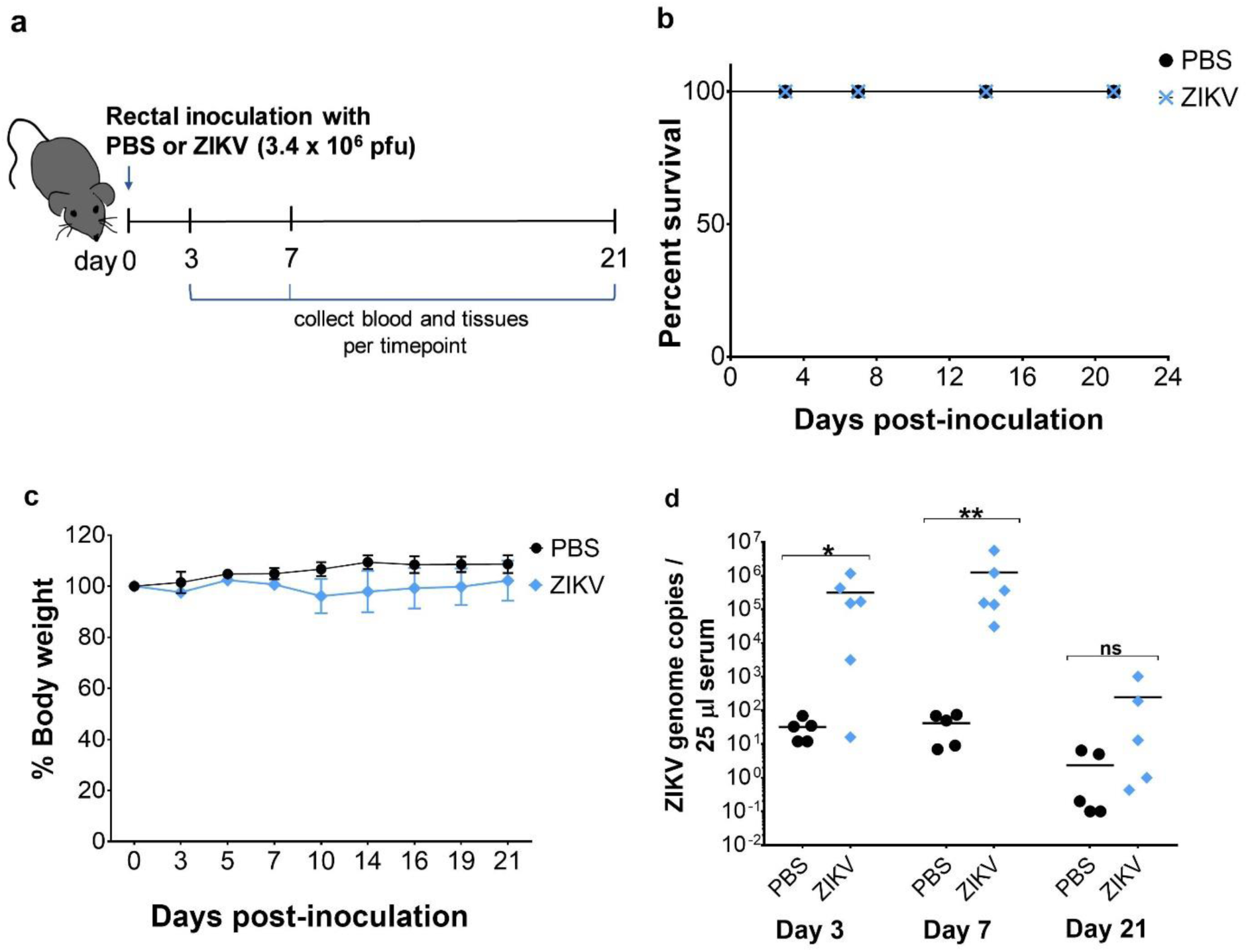
Rectal inoculation of ZIKV in *ifnar1*^*-/-*^ male mice results in a non-lethal, self-limiting infection with viremia. (A) Schematic showing the infection timeline for PBS (uninfected) and ZIKV infected groups. (B) Kaplan-Meier survival plot shows percent survival of *ifnar1*^-/-^ male mice post-rectal inoculation with PBS (n=5) or ZIKV (n=6). (C) Body weight change, represented as a percentage, of male mice post-rectal inoculation. Error bars represent the standard deviation (SD). (D) Viremia in uninfected and infected mice at 3, 7, and 21 dpi. Individual mice are represented by circles (PBS) or diamonds (ZIKV), with means represented by black lines. Two-tailed unpaired, non-parametric Mann-Whitney tests were conducted, where *p <0.05 and **p<0.001. ns, non-significant.

To assess ZIKV dissemination to the male reproductive system after rectal inoculation, we measured ZIKV genome copies in the testes and the accessory sex gland seminal vesicles, and the rectal mucosa at 3, 7 and 21 dpi by RT-qPCR. The rectal mucosa is lined by a single layer of columnar epithelial cells above the stroma that creates a mosaic of mucus-filled crypts. This simple columnar layer transitions into the stratified epithelium of the anal canal. We detected ZIKV replication within the rectal tissue at 3 dpi (Fig. 2A). ZIKV further disseminated to the testes and seminal vesicles, with peak acute viremic loads at 7 dpi (Fig. 2B), and persisted in the testes and rectal mucosa at 21 dpi (Fig. 2C). Future studies characterizing specific rectal cell types permissive to infection at earlier timepoints will provide insight on how ZIKV exploits this niche for replication and persistence.

**Figure 2.**
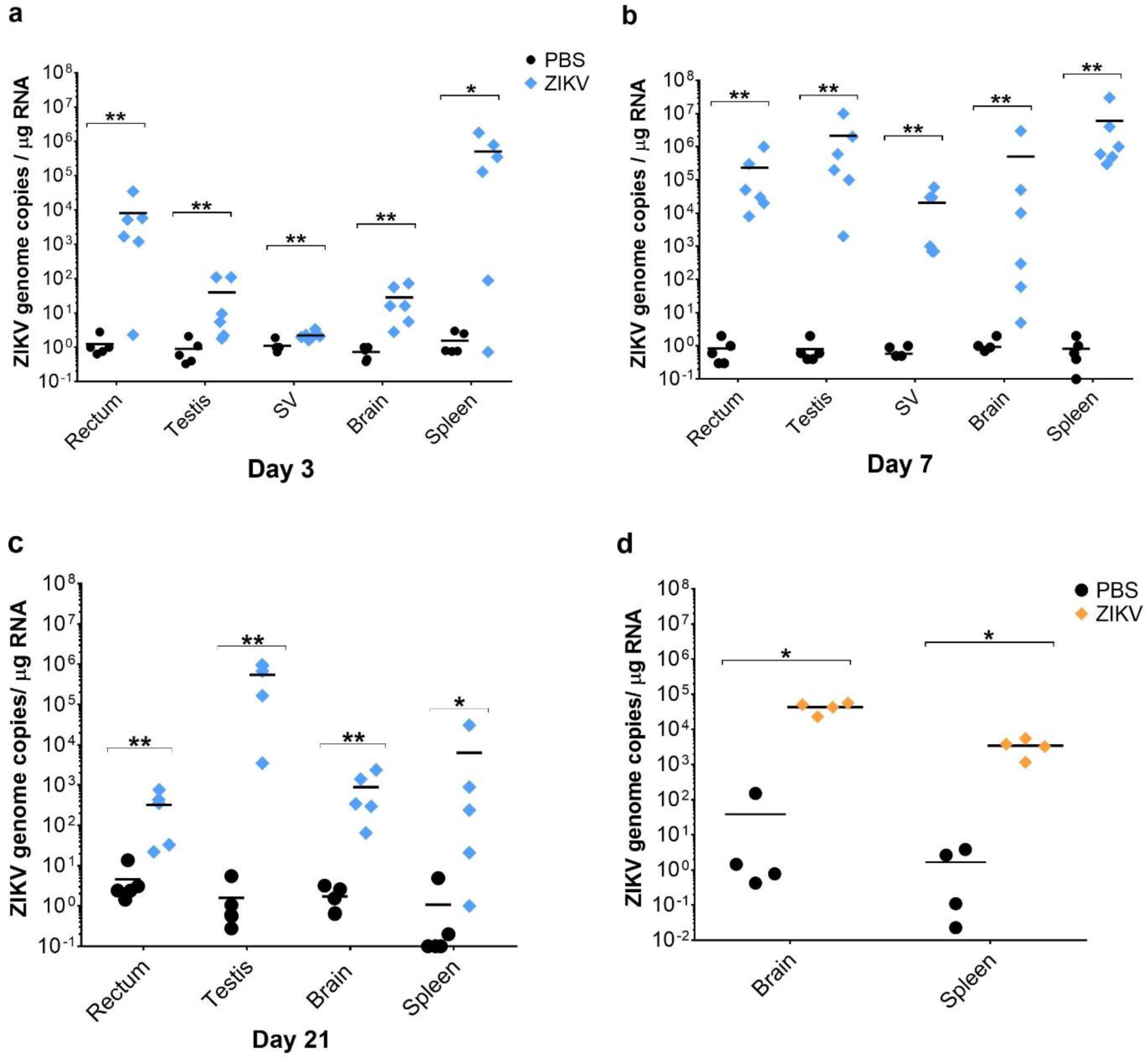
Tissue viral load following rectal and subcutaneous routes of ZIKV inoculation. Graphs depict ZIKV genome copies detected by RT-qPCR in various tissues at 3 dpi (A), 7 dpi (B), and 21 dpi (C). Each symbol corresponds to data from an individual mouse. n=5-6 mice per group. (D) Viral load in the spleen and brain at 6 days post-subcutaneous infection with ZIKV. n=4 mice per group. Two-tailed unpaired, non-parametric Mann-Whitney tests were conducted, where *p <0.05 and **p<0.001.

Rectal inoculation with ZIKV leads to viral dissemination to the brain at 3 dpi (Fig. 2A). Despite ZIKV infection of the brain, mice did not show malaise or neurological signs of disease at 7 dpi, as observed in subcutaneous route of infection. Some mice in the rectal route experiment showed little to no viral burden, while others sustained high viral loads in the brain (10^4^ -10^6^ ZIKV genome copies/μg of RNA). However, subcutaneously infected mice showed comparable higher viral loads in the brain at 6 dpi (10^5^ copies) (Fig. 2D), further suggesting that the virus was causing a fundamentally different disease outcome in rectally-infected mice. Furthermore, rectally-infected mice showed significantly high viral loads in the spleen at 3 dpi that persisted to 7 dpi (with a mean of 6×10^6^ copies) (Fig. 2A-B). In contrast, subcutaneously infected mice showed lower viral replication in the spleen (with ∼10^4^ copies per mouse) at 6 dpi (Fig. 2D). It is likely that during rectal infection, the virus can enter through the rectal mucosa and gain access to the inferior mesenteric vein (IMV) to reach the spleen. Before joining the hepatic portal vein, the IMV connects to the splenic vein and mixing of infected blood from the rectum to splenic circulation can occur. Subcutaneous inoculation with ZIKV can result in rapid entry of infectious viral particles into the circulatory system to reach the target organ brain.

We then determined whether *ifnar1*^*-/-*^ mice of reproductive age (12-week-old) would be able to control infection as efficiently as 4-6-week-old mice following rectal inoculation with ZIKV. 12-week-old mice did not succumb to infection nor showed signs of disease at 3 or 7 dpi (Supplementary Fig. 2). Infected mice were active and alert throughout the experimental period. Two of four infected mice showed relative body weight loss from day 5 to 7 and recovered by 10 dpi, with clearing of the viremia at 14 dpi (Supplementary Fig. 2B-C). Similar to the experimental results of young mice, ZIKV disseminated to the testes, seminal vesicles, and brain, and caused robust infection of the spleen by 14 dpi (Supplementary Fig. 2D). The lack of observable differences between age groups suggests that rectal inoculation with ZIKV may provide protective immunity that limits lethal infection as compared to subcutaneous route of infection, which induces acute encephalitis and neurological disease in this model system.

We then set out to investigate the cellular response to ZIKV infection in the spleens of 4-6-week-old mice in detail. Following rectal inoculation, ZIKV rapidly disseminated to the spleen at 3 dpi, further causing splenomegaly by 7 dpi (Fig. 3A-B). To determine the immune cells contributing to the splenic response to infection, we performed immunohistochemistry using markers for CD11b^+^ macrophages and dendritic cells, Ly6C^+^ neutrophils or granulocytes, and CD4^+^ T-cells. A rabbit polyclonal antibody recognizing ZIKV NS4B antigen was used to detect virus in the spleens of infected mice at 7 dpi. NS4B is a membrane bound cytoplasmic protein involved in flaviviral RNA genome replication. ZIKV infection caused infiltration or expansion of macrophages, dendritic cells, neutrophils, and CD4^+^ T-cells to the spleen (Fig. 3C). To identify ZIKV target cell types in the spleen, we performed co-immunostaining of ZIKV NS4B with cell-specific markers anti-CD4, anti-Ly6C, anti-CD11b, or anti-F4/80 on consecutive tissue sections. Viral antigen was not detected in CD4^+^ T-cells and neutrophils (Supplementary Fig. 3). Viral antigens were specifically found in the cytoplasm of F4/80^+^ macrophages in the red pulp, outside of germinal centers (Fig. 3D and Supplementary Fig. 4). Further studies examining virus-specific immune responses in the rectal mucosa and within peripheral organs, including lymph nodes and spleen, may provide insight on how the immune system controls ZIKV infection following rectal route of exposure.

**Figure 3.**
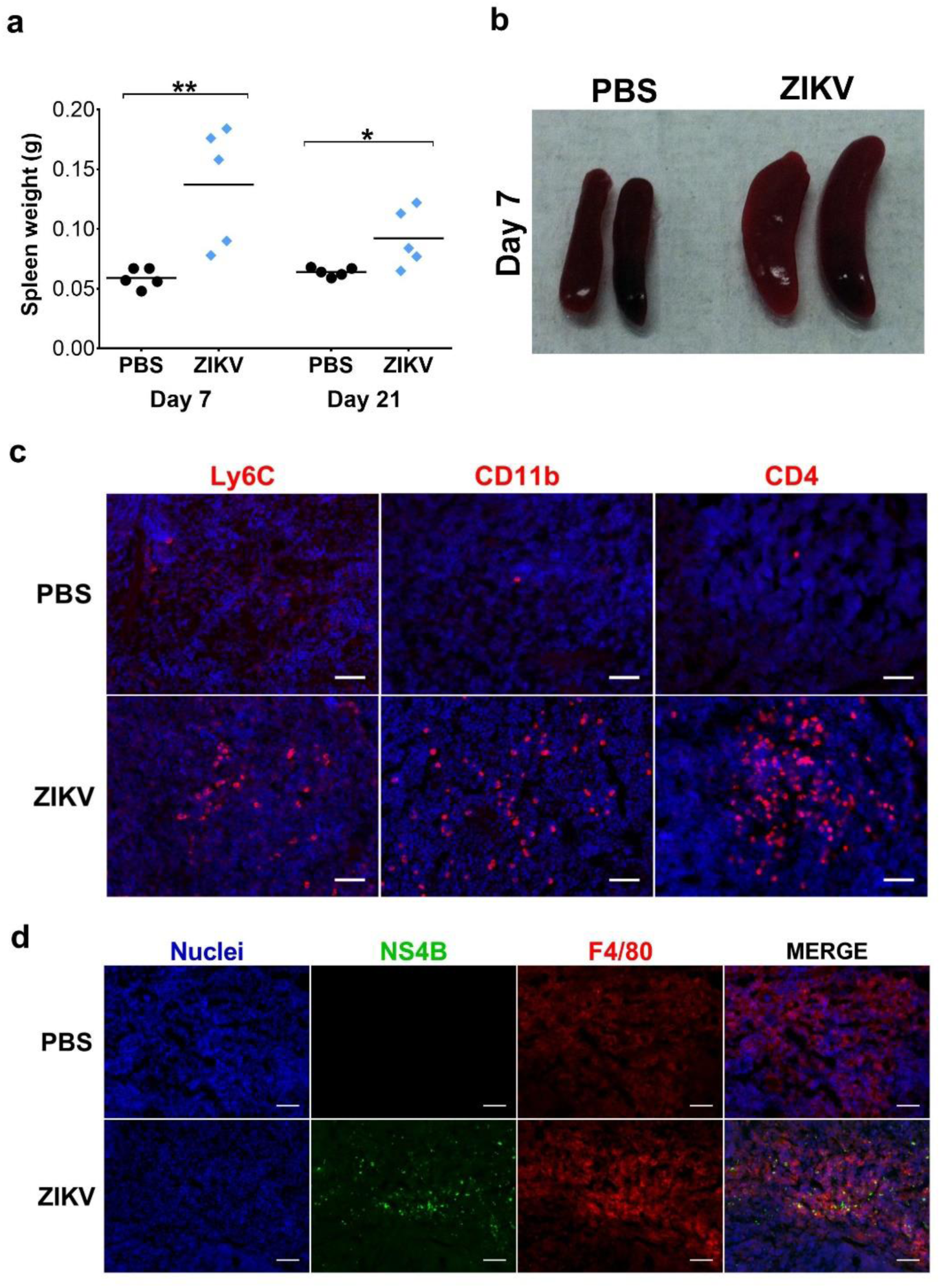
ZIKV induces splenomegaly and infiltration of inflammatory cells in spleen after rectal route of infection. (A) Graph shows spleen weight at 7 and 21 days post-rectal inoculation. (B) Gross image of spleens from PBS and ZIKV infected mice at 7 dpi. (C) Inflammatory and immune cellular infiltrates in the spleen of PBS and ZIKV inoculated mice at 7 dpi. Panels depict spleen tissue immunostained with anti-Ly6C, anti-CD11b, and anti-CD4. (D) Spleen tissue immunostained for ZIKV antigen NS4B. Anti-F4/80 was used to detect macrophages in the spleen. Merged images show viral antigen within the cytoplasm of F4/80^+^ macrophages in the red pulp of the spleen. Nuclei was stained with DAPI. Scale bar, 25 μm. The images (20X) are representative of 5-6 different animals for PBS and ZIKV groups.

**Figure 4.**
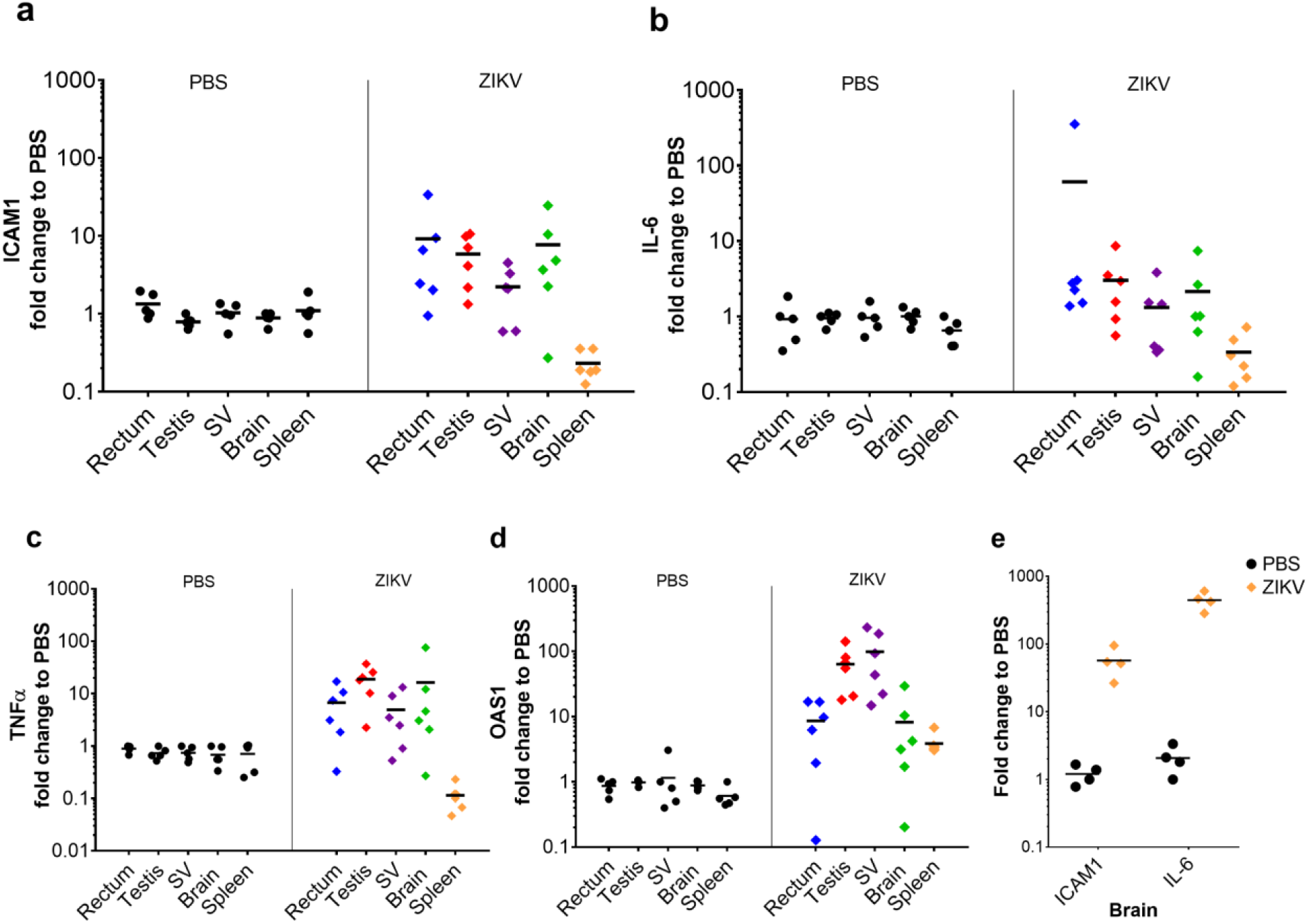
Tissue expression of inflammation-related genes at 7 days post-rectal inoculation with ZIKV. Expression of inflammation-related genes in the rectum, testes, seminal vesicles, brain, and spleen at 7 dpi. Total RNA was extracted from tissue and relative mRNA levels of ICAM1 (A), IL-6 (B), TNF-α (C), and OAS1 (D) were determined by RT-qPCR and normalized to GAPDH. The basal mRNA level in tissue from the PBS-inoculated group was normalized to 1 for each tissue type and the fold change relative to basal is shown. (E) Levels of ICAM1 and IL-6 in the brain of mice subcutaneously infected with ZIKV at 6 dpi.

During viral infection, there are active counter responses between the host and the virus, which often triggers the host’s first-line of antiviral defense, an innate immune response ^32^. This initial response is often measured by the induction of anti-viral and inflammatory cytokines at sites of active viral infection and tissue damage. Since the *ifnar1*^*-/-*^ mice contain intact NF-κB, as well as type II and III IFN signaling pathways, we observed that ZIKV elicited a robust innate immune response in the rectum, testes, seminal vesicles, and brain at 7 dpi, as provided by the activation of key inflammation-related genes (Fig. 4). The pro-inflammatory marker, ICAM1, was highly expressed in the rectum, testes, seminal vesicles, and brain, but not in the spleen (Fig. 4A). ZIKV also elicited a strong IL-6 (Fig. 4B) and TNFα (Fig. 4C) response at these sites. The interferon stimulated gene OAS1 was highly expressed in all the tissues tested, including the spleen (Fig. 4D), further suggesting a robust host response to infection mediated by an intrinsic cellular antiviral defense. During subcutaneous infection of ZIKV, systemic dissemination to the brain was not only confirmed by high viral load, but also by the vigorous IL-6 and ICAM1 response measured in the brains of infected mice (Fig. 4E). Overall, rectal inoculation of ZIKV prompts a robust initial innate immune response as observed by the expression of these pro-inflammatory and anti-viral cytokines, which can ultimately activate an anti-viral state to control infection. We expect that during rectal route of infection, ZIKV may stimulate stronger innate immune responses at the mucosal surface, which may further orchestrate the adaptive immune system to limit virus replication. This response can result in reduced tissue destruction and mortality.

Our current understanding of rectal route of ZIKV infection and transmission is limited. In the current study, we determined that the rectal mucosa is permissive to ZIKV replication following rectal route of inoculation, which may provide significant implications for ZIKV transmission via anorectal route among MSM and heterosexual partners. Our experiments with immunocompromised male mice show that ZIKV disseminates to the testes and seminal vesicles following rectal inoculation. The testes are considered an “immunoprivileged replication site” for ZIKV. Our study shows that ZIKV actively replicates in the testes at least 21 days post-infection, suggesting that the testes may provide a reservoir for active viral presence to perpetuate transmission. Semen, pre-ejaculate, or blood (via rectal damage or lacerations) may also contribute to ZIKV transmission via anorectal route. Therefore, studies examining the use of viricide to limit ZIKV entry to the rectal mucosa are critical. Moreover, rectal route of immunization against Rotavirus virus-like particles^33^ or a Hepatitis A virus vaccine^34^, have been shown to elicit strong systemic humoral responses. Our model system of rectal infection can be used as a platform to study immunization against ZIKV and to further understand the long-term effect of infection on mucosal immunity. It provides the opportunity for further exploring the impact of ZIKV persistence on male reproductive health and maternal transmission.

## Methods

### Ethics Statement

This study was performed in strict accordance with the recommendations of the Guide for the Care and Use of Laboratory Animal. The institutional Animal Care Use Committee of the Cedars-Sinai Medical Center approved the study.

### Cell lines and virus

*Aedes albopictus* mosquito C6/36 cell line (CRL-1660 cell line from ATCC) was used for propagating virus. C6/36 cells were maintained at 30°C in Dulbecco’s Modified Eagle’s Medium (Sigma Aldrich) supplemented with 10% heat-inactivated fetal bovine serum (FBS) (Gemini) with 100 U/mL penicillin (GIBCO) and 100 μg/mL streptomycin (GIBCO). Vero cells were obtained from ATCC and were cultured in DMEM supplemented with 10% FBS, penicillin and streptomycin, 1X GlutaMax (GIBCO), and 20 mM HEPES (GIBCO). The ZIKV PRVABC59 strain was received from the Center for Disease Control (CDC). Working ZIKV stocks were prepared by infecting C6/36 cells. Supernatants were collected and centrifuged at 200 g at 4°C for 10 min to remove cellular debris, aliquoted, and frozen at -80°C.

### Quantification of ZIKV by plaque assay

Monolayers of Vero cells were plated on 12-well plates. Virus supernatants from C6/36 cells were tittered by making dilutions in DMEM and were added to Vero cell monolayers at 37°C for 4 hours. Then, complete medium was added. Two days following the infection, plaques were counted manually as previously described^35^.

### Rectal inoculation of A129 mice

Pathogen-free *ifnar1* receptor-deficient mice (B6.129S2-*Ifnar1*^*tm1Agt*^/Mmjax or A129) were purchased from the Mutant Mouse Resource and Research Centers (MMRRC) supported by NIH. 4-6-week-old *ifnar1*^-/-^ male mice were rectally inoculated with PBS or ZIKV (PRVABC59, Puerto Rico 2015) while anesthetized. N=5-6 mice per group.

Briefly, a calginate swab was used to remove fecal stain from the anal orifice. For rectal ZIKV inoculation, each mouse was inoculated with 3.4×10^6^ plaque forming units (pfu) in a volume of 20 μl using a sterile smooth pipette tip (after passing or penetrating through anal orifice). Mice were kept with the rectum facing upwards for 4 minutes to reduce leakage of the inoculum. Survival, weight loss, and symptoms were monitored for 3, 7 and 21 days post-rectal inoculation (dpi). For the older mice study (12-week-old), rectal inoculated mice were infected for 14 days as described above. At the indicated post-infection times, mice were humanly euthanized using CO_2_ and blood was collected for viremia. To measure ZIKV viremia, 25 μl of blood was collected by direct cardiac puncture at the indicated timepoints post-infection. Total blood RNA was extracted using the viral RNA mini kit (Qiagen). The rectum, testes, seminal vesicles, spleen, and brain were harvested for evaluating viral load and cellular gene expression, as well as for immunohistochemistry. Tissues were removed from ZIKV infected mice or uninfected (PBS) control mice and dissected in thirds for fixation in 10% neutral buffered formalin, 4% paraformaldehyde, or RNAlater (Thermo Fisher Scientific) for RNA isolation.

### Subcutaneous inoculation of A129 mice

For subcutaneous (SC) infections with ZIKV, 4-6-week-old *ifnar1*^-/-^ male mice were inoculated with PBS (n=4 mice) or ZIKV (2 × 10^6^ pfu per mouse in a 40 μl volume) (n=4 mice) in the hind limb region. Blood was collected for viremia at 3 and 6 dpi, and the brain and spleen were harvested for viral load.

### RNA sample preparation and RT-qPCR

To determine levels of virus in tissues of control and infected mice, tissues were placed in RNAlater (Thermo Fisher Scientific) immediately after harvest and stored at -80°C. Tissues were homogenized and RNA was extracted using TRIzol as per the manufacturer (Thermo Fisher Scientific). Total RNA was isolated from rectum, testes, seminal vesicles, spleen, and brain. RNA was quantified using a NanoDrop 1000 Spectrophotometer (Thermo Fisher Scientific). cDNA was prepared from 1 μg of RNA using random hexamer primers and the SuperScript III Reverse Transcriptase Kit (Thermo Fischer Scientific). QPCR was performed using SYBR Green ROX Supermix (Life Technologies) and a QuantStudio 12K Flex Real-Time PCR System (ABI Thermo Fisher Scientific). Briefly, the amplification was performed using 10 μl volume reactions in a 384-well plate format with the following conditions: 50°C for 2 minutes followed by 95°C for 2 minutes, then 40 cycles at 95°C for 15 seconds and 60°C for 1 minute. The relative concentration of each transcript was calculated using the 2^-ΔCT^ method and Glyceraldehyde 3-phosphate dehydrogenase (GAPDH) threshold cycle (C_T_) values were used for normalization. The basal mRNA level in tissue from the PBS-inoculated group was normalized to 1 and the fold change relative to basal was determined for each tissue type. The qPCR primer pairs for the mRNA transcript targets are provided in Supplementary Table 1. ZIKV RNA transcript levels were quantified by comparing to a standard curve generated using dilutions (10^1^-10^9^ copies) of a ZIKV NS5 gene containing plasmid. ZIKV RNA levels are expressed as ZIKV genome copies per 1 μg of RNA using the standard curve.

### Immunohistochemistry

Tissue was incubated in 4% PFA for an hour and transferred to PBS. Tissues were then submerged in 10%, 20%, and 30% sucrose for an hour each. Tissue was then embedded in OCT (Fisher Healthcare) and incubated overnight at -80°C. Tissues were cut (10 μm thick) using a Leica cryostat microtome and mounted on Super Frost microscope slides (VWR). Sections were washed 3 times and permeabilized using blocking buffer (0.3% Triton X-100, 0.1% BSA, in 1 X PBS) for 1 hour at room temperature. For ZIKV staining, sections were incubated overnight at 4°C with a polyclonal anti-Zika NS4B antibody (rabbit, 1:250) (GeneTex). The sections were then rinsed with 1X PBS three times and incubated with secondary antibody, Alexa Fluor-conjugated 488 antibody (raised in rabbit; 1:1000), for 1 hour at room temperature. The antibodies used for immune cell staining are listed in Supplementary Table 2. For immune cell staining, Anti-Ly6C was used to immunostain monocytes, neutrophils, or granulocytes; anti-CD11b for monocytes, macrophages, and dendritic cells; anti-CD4 for CD4^+^ T-cells, and anti-F4/80 for macrophages. The secondary antibody, Alexa Fluor-conjugated 555 (1:1000), was used for visualization of immune cell infiltrates. Nucleus was stained with DAPI (4′,6-Diamidino-2-Phenylindole, Dihydrochloride) (Life Technologies) at a dilution of 1:1000 in blocking buffer.

### Data analysis

All testing was done at the two-sided alpha level of 0.05. Data were analyzed for statistical significance using the parametric two-tailed unpaired *t*-test Mann Whitney to compare two groups (uninfected vs. infected) with Graph Pad Prism software, version 7.0 (GraphPad Software, US). Kaplan-Meier survival curves were analyzed by the Gehan-Breslow-Wilcoxon test. *p*-values less than 0.05 were considered significant.

## Acknowledgments

We thank Leslie Garcia for assistance with tissue cryosectioning and preparation, and Preeti Hadavale and Constance Yue for help with immunohistochemistry. This research was funded by Cedars-Sinai Medical Center’s Institutional Research Award to V.A. and UCLA institutional support to R.S.

### Author contributions

L.E.M and V.A. designed the study. L.E.M, G.G, D.C, and V.A. performed experiments. L.E.M. and D.C. performed data analysis and interpretation. L.E.M, G.G., D.C., and V.A. wrote the manuscript.

### Additional information

Supplementary information is available for this paper. Correspondence and requests for materials should be addressed to V.A.

### Competing interests

The authors declare no competing financial interests.

## Supplementary Information

**Supplementary Figure 1.**
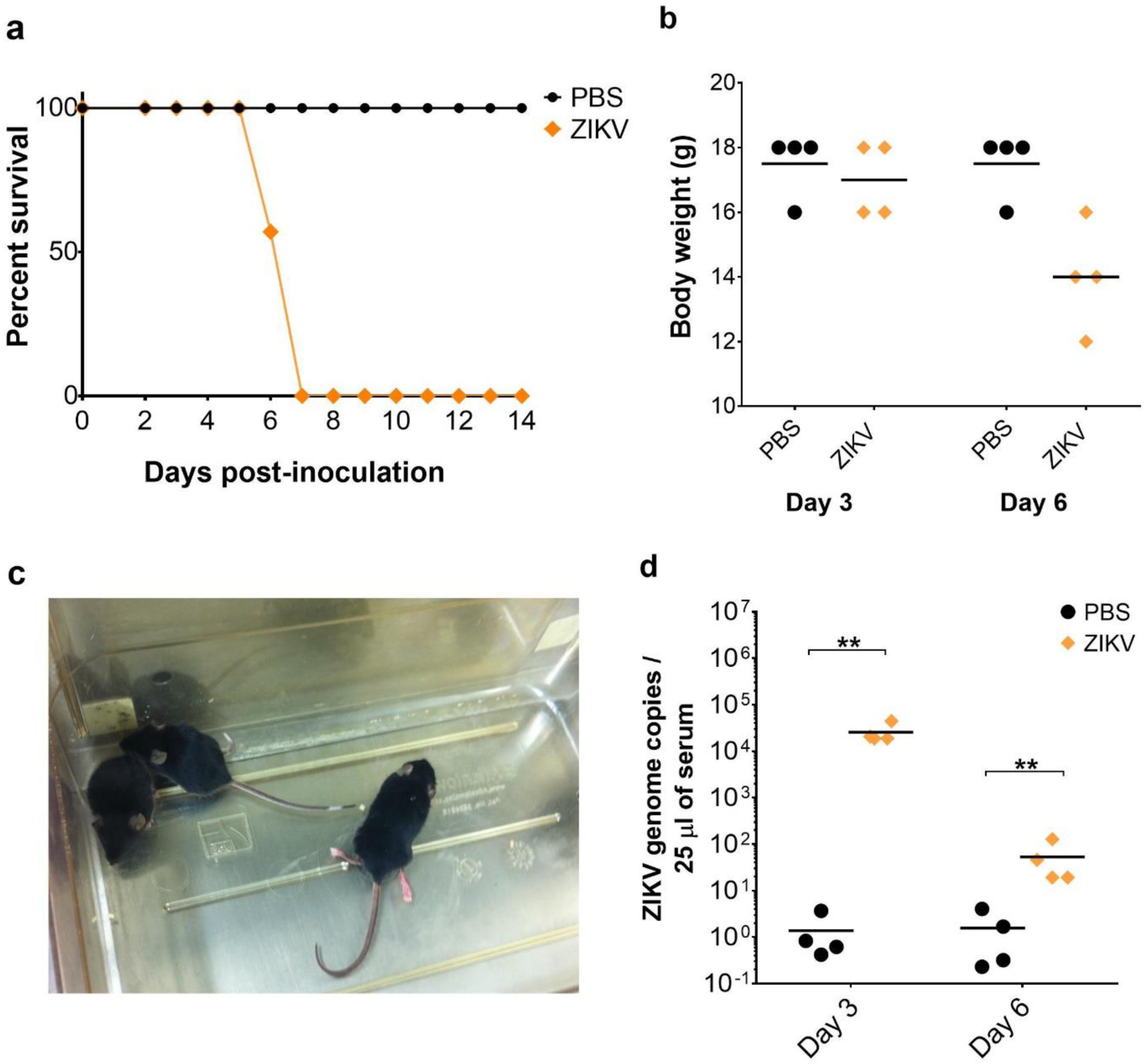

**Supplementary Figure 2.**
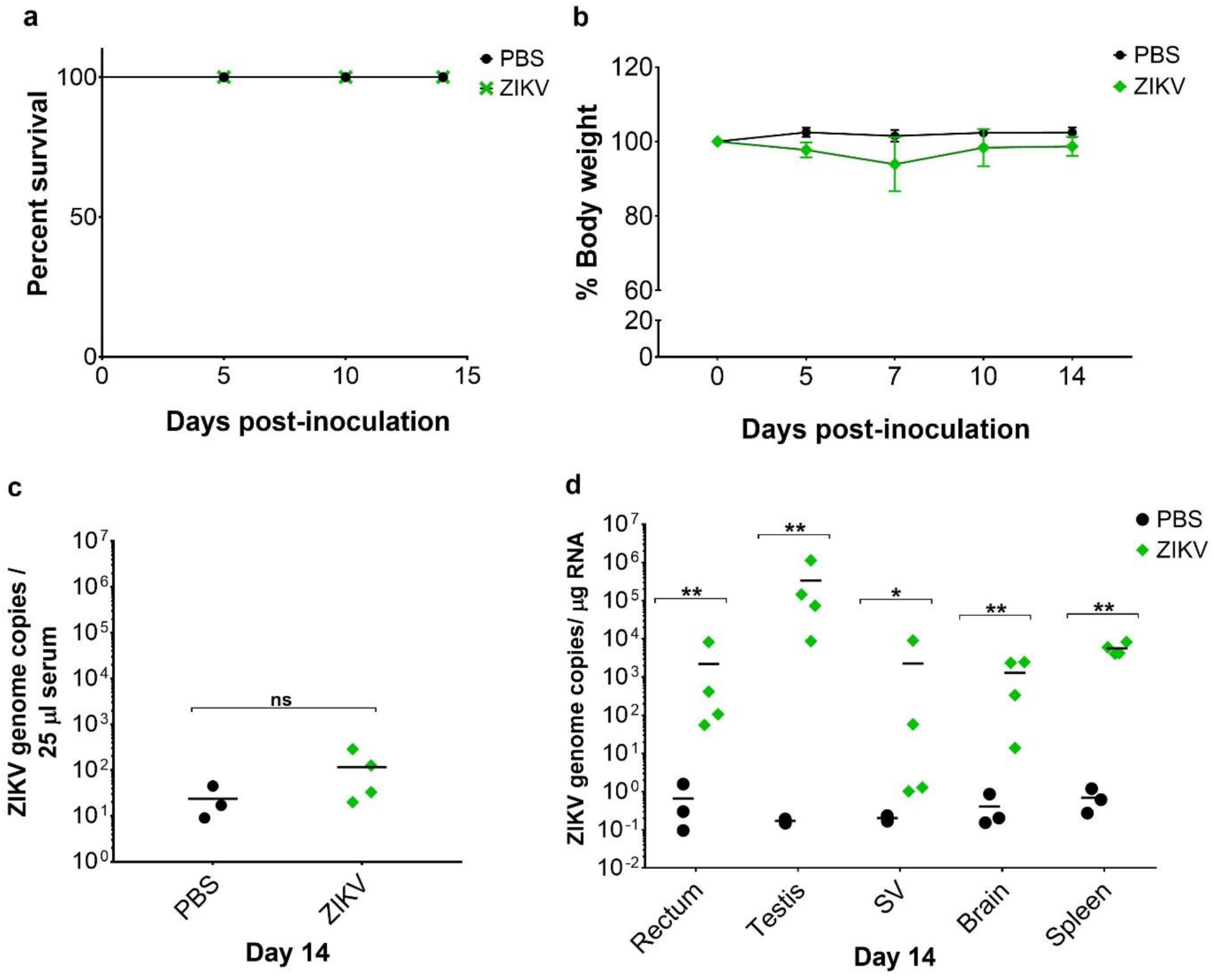

**Supplementary Figure 3.**
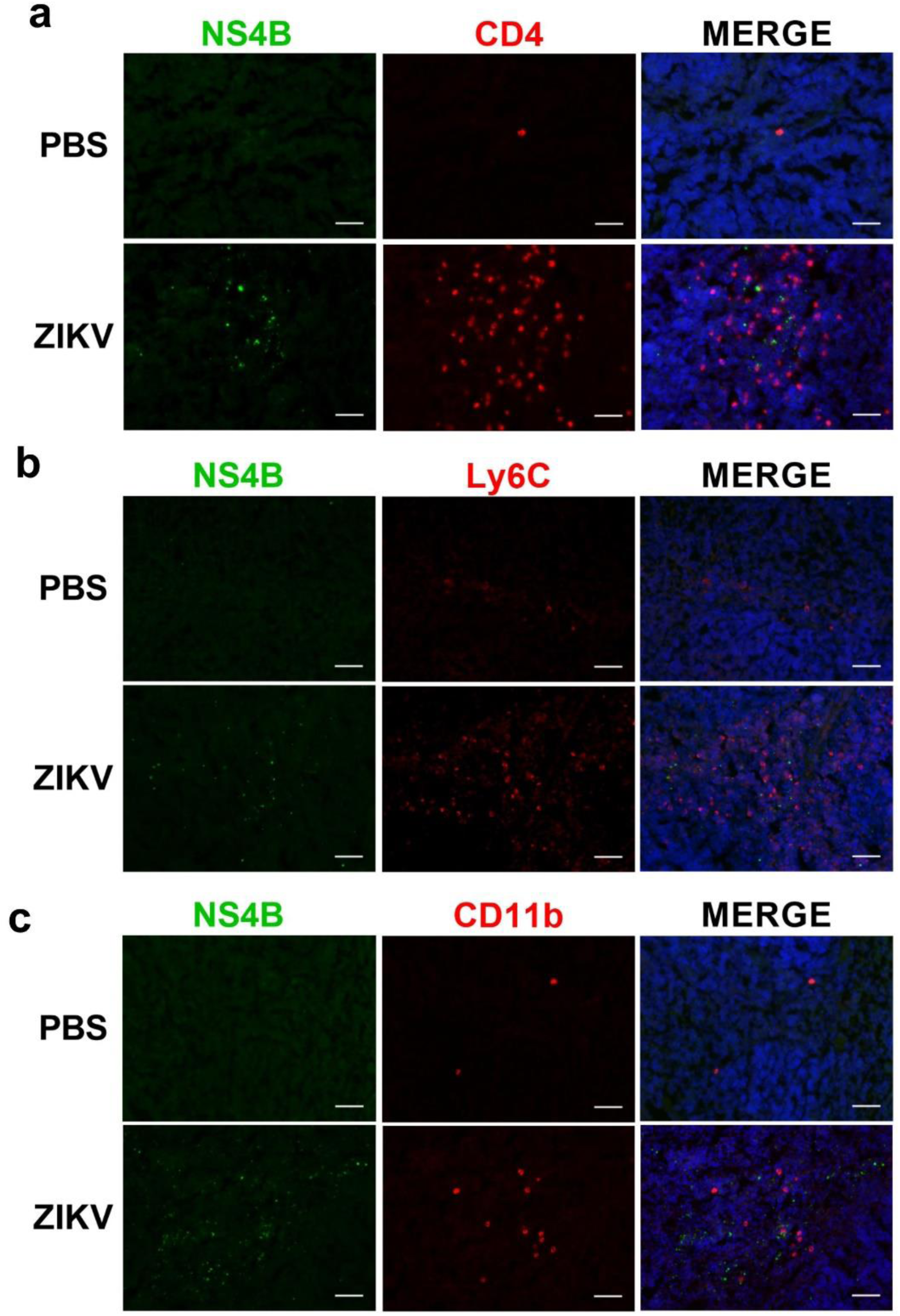

**Supplementary Figure 4.**
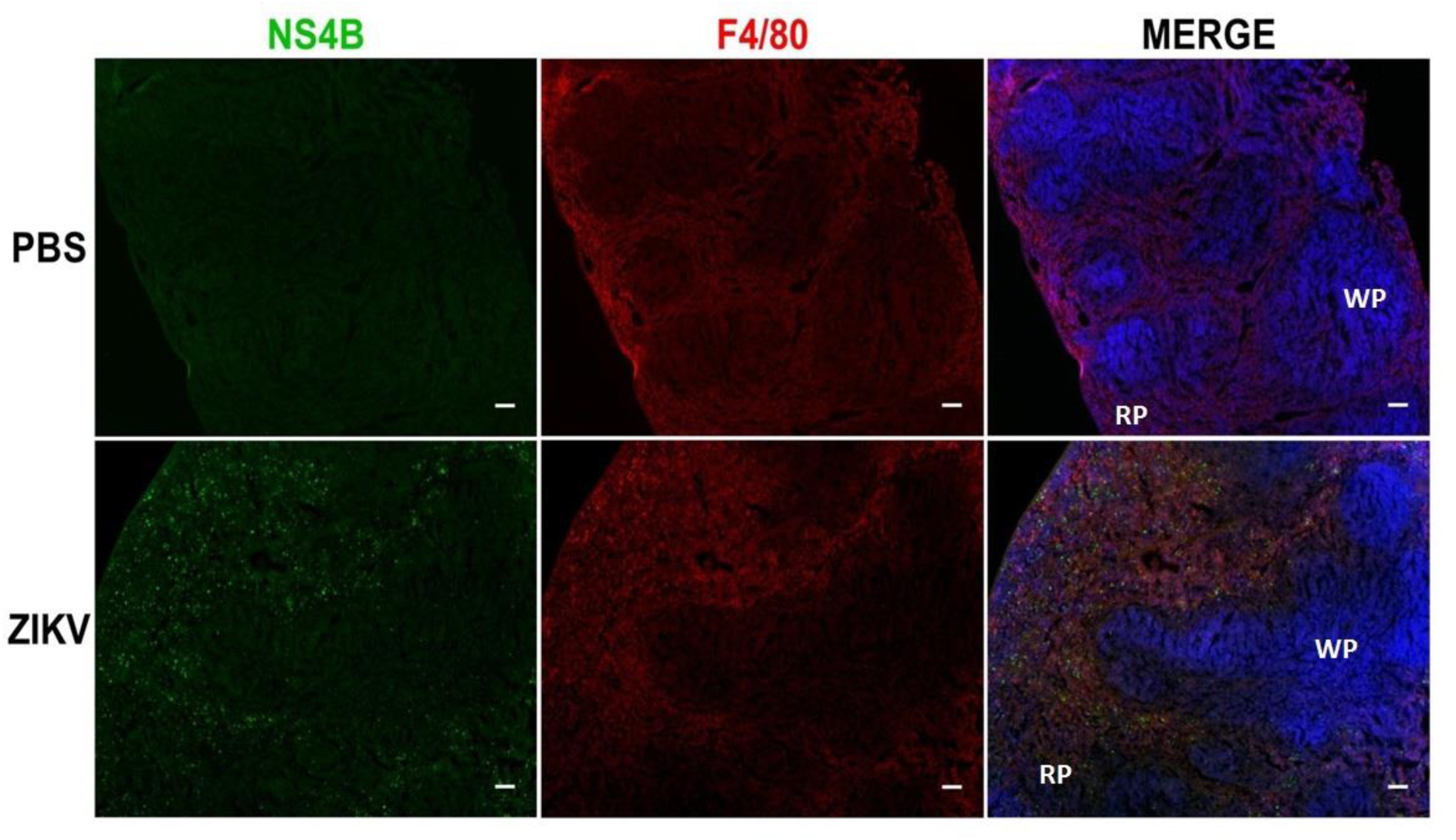

**Supplementary Table 1.**
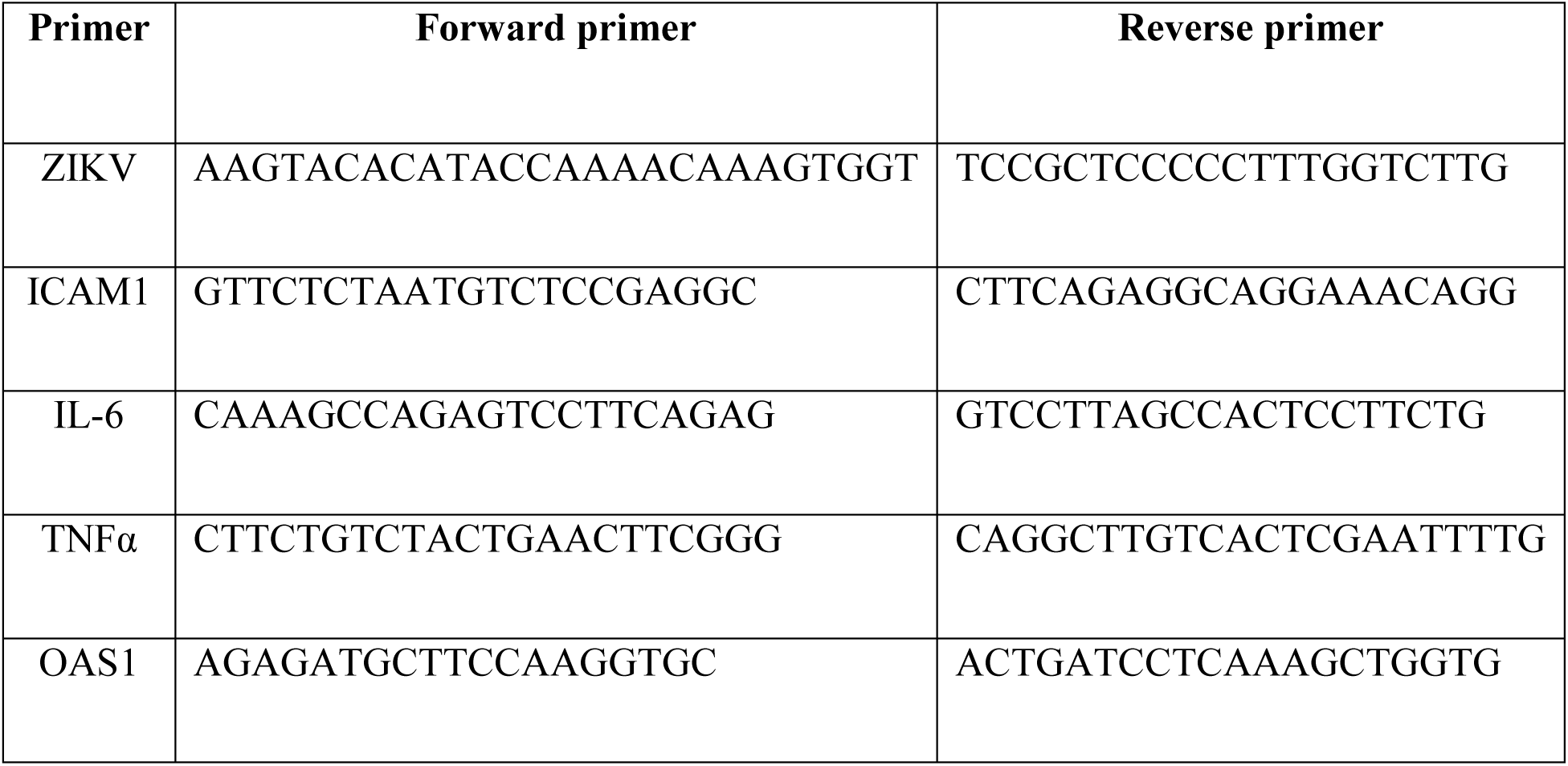
RT-qPCR primer sequences

**Supplementary Table 2.**
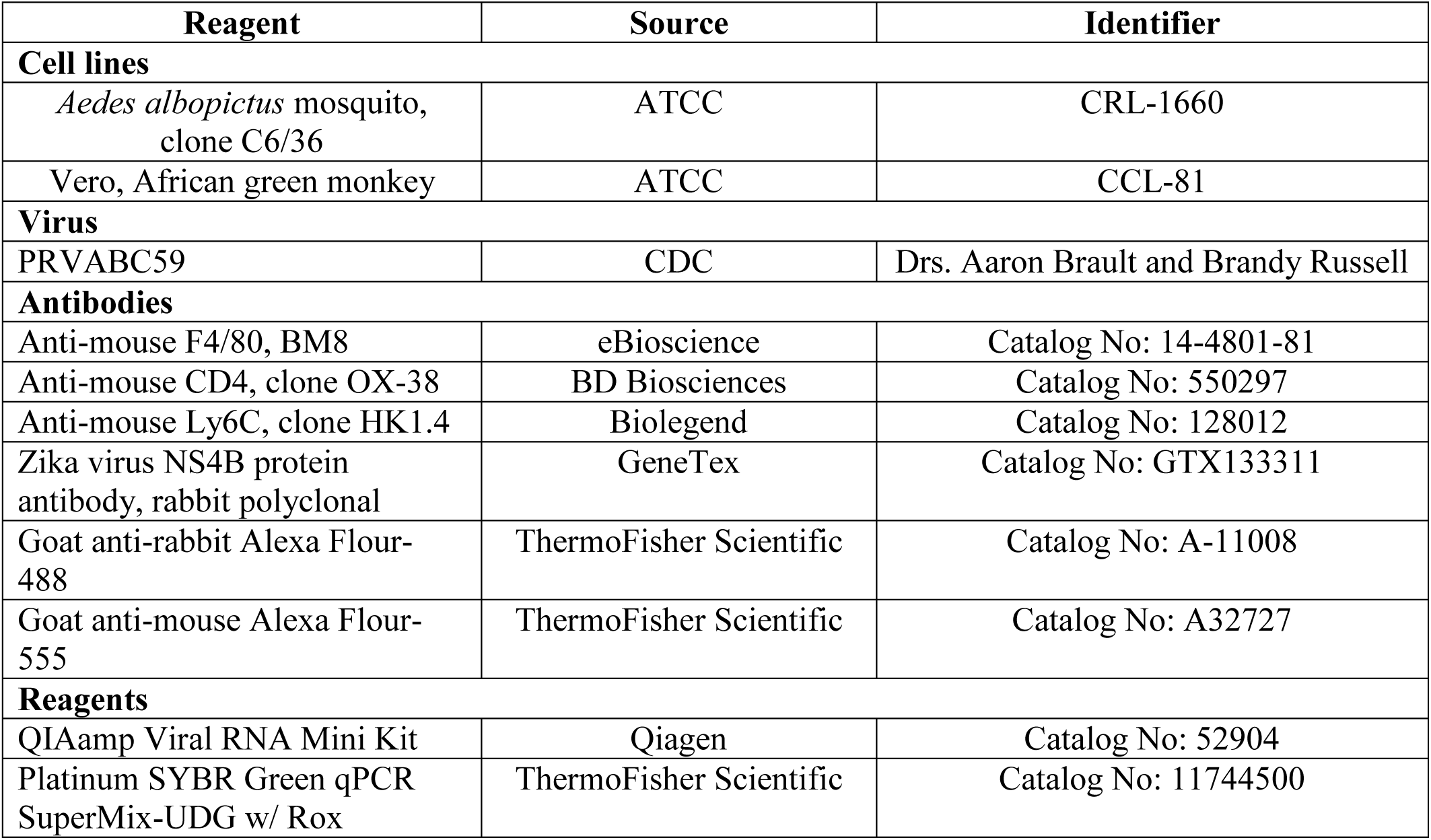
Reagents or resources used in this study

| Reagent | Source | Identifier |
| --- | --- | --- |
| **Cell lines** |  |  |
| *Aedes albopictus* mosquito, clone C6/36 | ATCC | CRL-1660 |
| Vero, African green monkey | ATCC | CCL-81 |
| **Virus** |  |  |
| PRVABC59 | CDC | Drs. Aaron Brault and Brandy Russell |
| **Antibodies** |  |  |
| Anti-mouse F4/80, BM8 | eBioscience | Catalog No: 14-4801-81 |
| Anti-mouse CD4, clone OX-38 | BD Biosciences | Catalog No: 550297 |
| Anti-mouse Ly6C, clone HK1.4 | Biolegend | Catalog No: 128012 |
| Zika virus NS4B protein antibody, rabbit polyclonal | GeneTex | Catalog No: GTX133311 |
| Goat anti-rabbit Alexa Flour-488 | ThermoFisher Scientific | Catalog No: A-11008 |
| Goat anti-mouse Alexa Flour-555 | ThermoFisher Scientific | Catalog No: A32727 |
| **Reagents** |  |  |
| QIAamp Viral RNA Mini Kit | Qiagen | Catalog No: 52904 |
| Platinum SYBR Green qPCR SuperMix-UDG w/ Rox | ThermoFisher Scientific | Catalog No: 11744500 |

**Supplementary Figure 1. Subcutaneous infection with ZIKV causes weight loss, posterior hind limb paralysis, and mortality in *ifnar1*^*-/-*^ mice.** *Related to Figure 1.* (A) Kaplan-Meier survival plot and (B) body weight change of PBS and ZIKV-infected mice via subcutaneous route. (C) A mouse (right) in the infected group shows hind limb posterior paralysis at 6 dpi. (D) Viremia at 3 and 6 dpi. Two-tailed unpaired, non-parametric Mann-Whitney tests were conducted, where *p <0.05 and **p<0.001.

**Supplementary Figure 2. ZIKV burden at 14 days post-rectal inoculation of 12-week-old male mice.** *Related to Figures 1 and 2.* (A) Kaplan-Meier survival plot of PBS (n=3) and ZIKV (n=4) infected groups. (B) Graph presents percent body weight change at various timepoints post-rectal inoculation. (C) Viremia in PBS and ZIKV-infected mice at 14 dpi. (D) Tissue viral load at 14 dpi. Each symbol corresponds to data from an individual mouse. Two-tailed unpaired, student t-tests were conducted, where *p <0.05 and **p<0.001**.

**Supplementary Figure 3. ZIKV antigens were not present in immune cells of the spleen.** *Related to Figure 3C.* Uninfected and ZIKV infected spleen tissue at 7 dpi were co-immunostained with ZIKV NS4B and anti-CD4 (A), anti-Ly6C (B), or anti-CD11b (C). Nuclei was stained with DAPI. Scale bar, 25 μm. The images (20X) are representative of 5 different mice for PBS and 6 mice for the ZIKV infected group.

**Supplementary Figure 4. ZIKV antigens localize with splenic macrophages within the red pulp of the spleen after rectal route of infection.** *Related to Figure 3D.* Uninfected and ZIKV infected spleen at 7 dpi were immunostained for ZIKV protein NS4B and macrophages using anti-F4/80 antibody. Merged images show ZIKV antigens present in the cytoplasm of F4/80^+^ macrophages within the red pulp (RP) of the spleen. White pulp, WP. Nuclei was stained with DAPI. Scale bar, 100 μm. The images (4X) are representative of 5 different mice for PBS and 6 mice for the ZIKV infected group.

## References

1 Cao-Lormeau, V. M. et al. Zika virus, French polynesia, South pacific, 2013. Emerg Infect Dis 20, 1085–1086, doi:10.3201/eid2006.140138 (2014).

2 Ventura, C. V., Maia, M., Bravo-Filho, V., Gois, A. L. & Belfort, R., Jr. Zika virus in Brazil and macular atrophy in a child with microcephaly. Lancet 387, 228, doi:10.1016/S0140-6736(16)00006-4 (2016).

3 Ventura, C. V., Maia, M., Dias, N., Ventura, L. O. & Belfort, R., Jr. Zika: neurological and ocular findings in infant without microcephaly. Lancet 387, 2502, doi:10.1016/S0140- 6736(16)30776-0 (2016).

4 Ventura, C. V. et al. Risk Factors Associated With the Ophthalmoscopic Findings Identified in Infants With Presumed Zika Virus Congenital Infection. JAMA Ophthalmol 134, 912–918, doi:10.1001/jamaophthalmol.2016.1784 (2016).

5 Ventura, C. V. et al. Optical Coherence Tomography of Retinal Lesions in Infants With Congenital Zika Syndrome. JAMA Ophthalmol 134, 1420–1427, doi:10.1001/jamaophthalmol.2016.4283 (2016).

6 Hills, S. L. et al. Transmission of Zika Virus Through Sexual Contact with Travelers to Areas of Ongoing Transmission - Continental United States, 2016. MMWR Morb Mortal Wkly Rep 65, 215–216, doi:10.15585/mmwr.mm6508e2 (2016).

7 Moreira, J., Peixoto, T. M., Machado de Siqueira, A. & Lamas, C. C. Sexually acquired Zika virus: a systematic review. Clin Microbiol Infect, doi:10.1016/j.cmi.2016.12.027 (2017).

8 Russell, K. et al. Male-to-Female Sexual Transmission of Zika Virus-United States, January-April 2016. Clin Infect Dis 64, 211–213, doi:10.1093/cid/ciw692 (2017).

9 Deckard, D. T. et al. Male-to-Male Sexual Transmission of Zika Virus--Texas, January 2016. MMWR Morb Mortal Wkly Rep 65, 372–374, doi:10.15585/mmwr.mm6514a3 (2016).

10 Davidson, A., Slavinski, S., Komoto, K., Rakeman, J. & Weiss, D. Suspected Female-to-Male Sexual Transmission of Zika Virus - New York City, 2016. MMWR Morb Mortal Wkly Rep 65, 716–717, doi:10.15585/mmwr.mm6528e2 (2016).

11 Atkinson, B. et al. Detection of Zika Virus in Semen. Emerg Infect Dis 22, 940, doi:10.3201/eid2205.160107 (2016).

12 Gaskell, K. M., Houlihan, C., Nastouli, E. & Checkley, A. M. Persistent Zika Virus Detection in Semen in a Traveler Returning to the United Kingdom from Brazil, 2016. Emerg Infect Dis 23, 137–139, doi:10.3201/eid2301.161300 (2017).

13 Harrower, J. et al. Sexual Transmission of Zika Virus and Persistence in Semen, New Zealand, 2016. Emerg Infect Dis 22, 1855–1857, doi:10.3201/eid2210.160951 (2016).

14 Mansuy, J. M. et al. Zika virus: high infectious viral load in semen, a new sexually transmitted pathogen? Lancet Infect Dis 16, 405, doi:10.1016/S1473-3099(16)00138-9 (2016).

15 Mansuy, J. M. et al. Zika virus in semen of a patient returning from a non-epidemic area. Lancet Infect Dis 16, 894–895, doi:10.1016/S1473-3099(16)30153-0 (2016).

16 Matheron, S. et al. Long-Lasting Persistence of Zika Virus in Semen. Clin Infect Dis 63, 1264, doi:10.1093/cid/ciw509 (2016).

17 Atkinson, B. et al. Presence and Persistence of Zika Virus RNA in Semen, United Kingdom, 2016. Emerg Infect Dis 23, doi:10.3201/eid2304.161692 (2017).

18 Nicastri, E. et al. Persistent detection of Zika virus RNA in semen for six months after symptom onset in a traveller returning from Haiti to Italy, February 2016. Euro Surveill 21, doi:10.2807/1560-7917.ES.2016.21.32.30314 (2016).

19 Oliveira Souto, I. et al. Persistence of Zika virus in semen 93 days after the onset of symptoms. Enferm Infecc Microbiol Clin, doi:10.1016/j.eimc.2016.10.009 (2016).

20 Catania, J. A. et al. The continuing HIV epidemic among men who have sex with men. Am J Public Health 91, 907–914 (2001).

21 Aliota, M. T. et al. Characterization of Lethal Zika Virus Infection in AG129 Mice. PLoS Negl Trop Dis 10, e0004682, doi:10.1371/journal.pntd.0004682 (2016).

22 Lazear, H. M. et al. A Mouse Model of Zika Virus Pathogenesis. Cell Host Microbe 19, 720–730, doi:10.1016/j.chom.2016.03.010 (2016).

23 Miner, J. J. et al. Zika Virus Infection during Pregnancy in Mice Causes Placental Damage and Fetal Demise. Cell 165, 1081–1091, doi:10.1016/j.cell.2016.05.008 (2016).

24 Rossi, S. L. et al. Characterization of a Novel Murine Model to Study Zika Virus. Am J Trop Med Hyg 94, 1362–1369, doi:10.4269/ajtmh.16-0111 (2016).

25 Khan, S. et al. Dampened antiviral immunity to intravaginal exposure to RNA viral pathogens allows enhanced viral replication. J Exp Med 213, 2913–2929, doi:10.1084/jem.20161289 (2016).

26 Yockey, L. J. et al. Vaginal Exposure to Zika Virus during Pregnancy Leads to Fetal Brain Infection. Cell 166, 1247–1256 e1244, doi:10.1016/j.cell.2016.08.004 (2016).

27 Govero, J. et al. Zika virus infection damages the testes in mice. Nature 540, 438–442, doi:10.1038/nature20556 (2016).

28 Uraki, R. et al. Zika virus causes testicular atrophy. Sci Adv 3, e1602899, doi:10.1126/sciadv.1602899 (2017).

29 Ma, W. et al. Zika Virus Causes Testis Damage and Leads to Male Infertility in Mice. Cell 168, 542, doi:10.1016/j.cell.2017.01.009 (2017).

30 Duggal, N. K. et al. Frequent Zika Virus Sexual Transmission and Prolonged Viral RNA Shedding in an Immunodeficient Mouse Model. Cell Rep 18, 1751–1760, doi:10.1016/j.celrep.2017.01.056 (2017).

31 Manangeeswaran, M., Ireland, D. D. & Verthelyi, D. Zika (PRVABC59) Infection Is Associated with T cell Infiltration and Neurodegeneration in CNS of Immunocompetent Neonatal C57Bl/6 Mice. PLoS Pathog 12, e1006004, doi:10.1371/journal.ppat.1006004 (2016).

32 Katze, M. G., Fornek, J. L., Palermo, R. E., Walters, K. A. & Korth, M. J. Innate immune modulation by RNA viruses: emerging insights from functional genomics. Nat Rev Immunol 8, 644–654, doi:10.1038/nri2377 (2008).

33 Parez, N. et al. Rectal immunization with rotavirus virus-like particles induces systemic and mucosal humoral immune responses and protects mice against rotavirus infection. J Virol 80, 1752–1761, doi:10.1128/JVI.80.4.1752-1761.2006 (2006).

34 Mitchell, L. A. & Galun, E. Rectal immunization of mice with hepatitis A vaccine induces stronger systemic and local immune responses than parenteral immunization. Vaccine 21, 1527–1538 (2003).

35 Contreras, D. & Arumugaswami, V. Zika Virus Infectious Cell Culture System and the In Vitro Prophylactic Effect of Interferons. J Vis Exp, doi:10.3791/54767 (2016).

